# Identifying relationships between imaging phenotypes and lung cancer-related mutation status: *EGFR* and *KRAS*

**DOI:** 10.1101/794123

**Authors:** Gil Pinheiro, Tania Pereira, Catarina Dias, Cláudia Freitas, Venceslau Hespanhol, José Luis Costa, António Cunha, Hélder P. Oliveira

## Abstract

*EGFR* and *KRAS* are the most frequently mutated genes in lung cancer, being active research topics in targeted therapy. Biopsy is the traditional method to genetically characterise a tumour. However, it is a risky procedure, painful for the patient, and, occasionally, the tumour might be inaccessible. This work aims to study and debate the nature of the relationships between imaging phenotypes and lung cancer-related mutation status. Until now, the literature has failed to point to new research directions, mainly consisting of results-oriented works in a field where there is still not enough available data to train clinically viable models. We intend to open a discussion about critical points and to present new possibilities for future radiogenomics studies. We conducted high-dimensional data visualisation and developed classifiers, which allowed us to analyse the results for *EGFR* and *KRAS* biological markers according to different combinations of input features. We show that *EGFR* mutation status might be correlated to CT scans imaging phenotypes, however, the same does not seem to hold true for *KRAS* mutation status. Also, the experiments suggest that the best way to approach this problem is by combining nodule-related features with features from other lung structures.

## Introduction

Lung cancer is the cancer type leading the incidence and mortality rates ^1, 2^. This is linked to the fact that it is often diagnosed in an advanced stage, with 15% or less chance of a 5-year survival^3^, which magnifies the importance of treatments for advanced-stage disease. In Non-small-cell lung cancer (NSCLC), which constitutes 85% of all cases of lung cancer^4^, certain genomic biomarkers are now considered critical for the prognostic. By determining their mutation status, it is possible to provide a targeted therapy for each patient^5^. Epidermal Growth Factor Receptor (*EGFR*) and Kristen Rat Sarcoma Viral Oncogene Homolog (*KRAS*) are the most frequently mutated gene in lung cancer^6^. *EGFR* is over-expressed in 40 to 80% of NSCLC patients from never-smokers and it is associated with poor diagnosis^6, 7^. Unlike the previous marker, *KRAS* is associated with tobacco use, with only 5 to 10% of *KRAS*-mutant lung cancers arising in never or light smokers^6, 8^. Current molecularly-targeted therapies can effectively target specific biomarkers, decreasing multiple undesirable side effects associated with cancer treatment^9^. *EGFR* mutations are the most well characterised and several NSCLC treatments based on it were already approved^9^. Effective therapies targeting *KRAS* are yet to be developed, although it is an extremely relevant topic that is actively being researched^10^.

The traditional method of characterising the tumour is by extracting tumour tissue in a biopsy, which is then analysed through different approaches, such as genomic-based ones^11^. In spite of being a successful approach in clinical oncology, biopsies tend to increase medical complications. Thus, there is the urge to find a less invasive way to shape the treatment^12^. Medical image analysis can help to solve these issues in two ways: by providing tools capable of measuring characteristics of the lung and, more specifically, the tumour; and with models that use only image features to obtain results through automatic or semi-automatic processes. These models can either use qualitative features, obtained by semantic annotations from human observers, or use quantitative features, obtained through a radiomic approach, which extracts features directly from the image^13^. Radiogenomics, a specific field within radiomics, is defined by the correlation between quantitative features, directly extracted from radiological images (imaging phenotype), and genetic information (genotype)^14^. Studies in lung cancer have presented the association between *EGFR* mutation status and quantitative features extracted from computed tomography (CT) scans^14–17^. The most recent methods are based on convolutional neural networks, which are end-to-end approaches that allow to automatically learn the whole process, reducing the subjectivity and human effort^14, 18^. Also, regarding qualitative features, recent works have shown that semantic human annotations of CT scans can be used to train a model to accurately predict *EGFR* mutation status, although the same was not verifiable for *KRAS*^19, 20^.

Our previous work was a preliminary study which used a public database^13, 21, 22^ to create predictive models for *EGFR* and *KRAS*^20^. In the current work, we apply a more robust approach based on multiple splits to define the train and test sets, preventing an eventual bias from specific patients and effectively assessing the variance in the data. Additionally, this study aims to provide further advances and to open new discussions in the lung cancer radiogenomics field by exploring the data and building machine learning models, while considering different subsets of inputs. More specifically, predictive models for *EGFR* and *KRAS* mutation status in lung cancer were developed. Following the current direction in the literature, where the analysis only focuses on the nodule structure and texture^23, 24^, we started by using objective radiomic features directly extracted from nodules in CT scans. Then, semantic features, annotated during radiologist evaluation, were used as input. Unlike their radiomic counterpart, they comprise not only nodule characteristics, but also lung characteristics external to the tumour. Clinical features as patient’s gender and smoking status were considered due to its significant association with mutation status prevalence, confirmed in recent studies^25–27^. Moreover, the comparison between its results and those found in the literature review is also presented and is suggested a new perspective about gene mutation status prediction based on image analysis.

## Results

### Data Visualisation

When using Principal Component Analysis (PCA) followed by t-distributed Stochastic Neighbour Embedding (t-SNE) for dimensionality reduction, it is possible to conclude that the separation of classes between mutated and wild type *EGFR* gene status is better when using hybrid semantic features. However, the separation is not perfect, as there are samples outside their cluster, which illustrates the level of complexity faced in a classification process (Figure 1a). Contrarily, for *KRAS*, there is no visible separation between classes with any type of input features (Figure 1b). The remaining data visualisation images can be found in Supplementary Figure 1.

**Figure 1.**
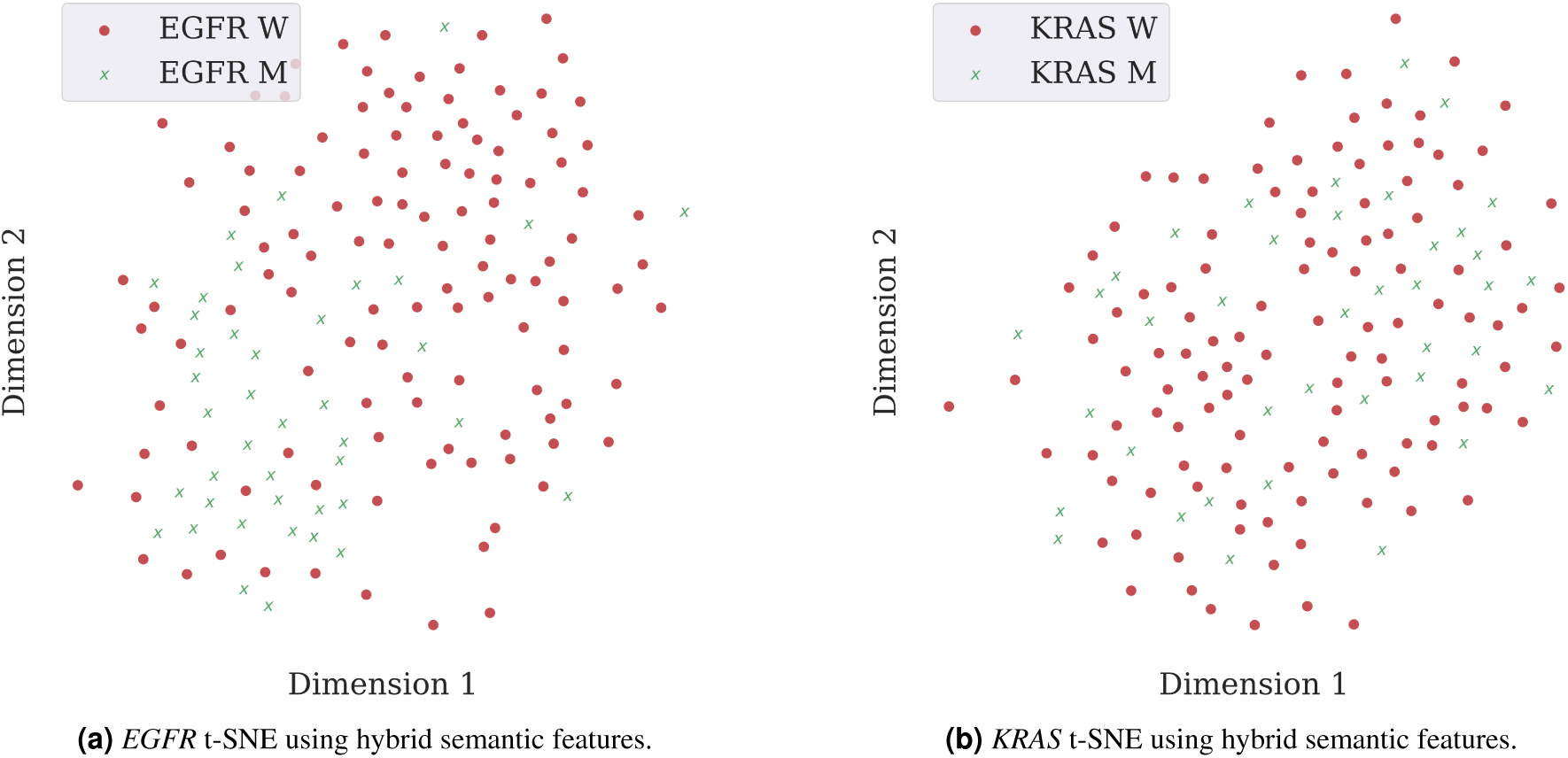
Visualisation of sample distributions based on PCA and t-SNE. Each point is coloured according to its mutation status, with red dots and green crosses representing wild type and mutated cases, respectively.

### Classification Results

Mean values of Area Under the Curve (AUC) were reported for 100 random data splits, with a division of 80% and 20% for training and testing, respectively. Two main types of input features were considered: radiomic and semantic. The semantic were further divided into features that only describe the nodule, features that only describe structures external to nodule and a hybrid between the previous two. Radiomics were not further divided as they only describe the nodule.

Only the predictive models for *EGFR* showed relevant results, with a maximum mean AUC of 0.7458 ± 0.0877 using the hybrid semantic features (represented by the mean ROC curve in Figure 2). The second best result was obtained using non-nodule semantic features. The worst results were obtained using features only from the nodule, using radiomic and semantic type of features. For *KRAS*, it was not possible to build any acceptable model. The performance results confirm the difficulty of gene mutation status classification, which is visible in the t-SNE projections, where there was not possible to achieve a clear separation between classes (Figure 1).

**Table 1.**
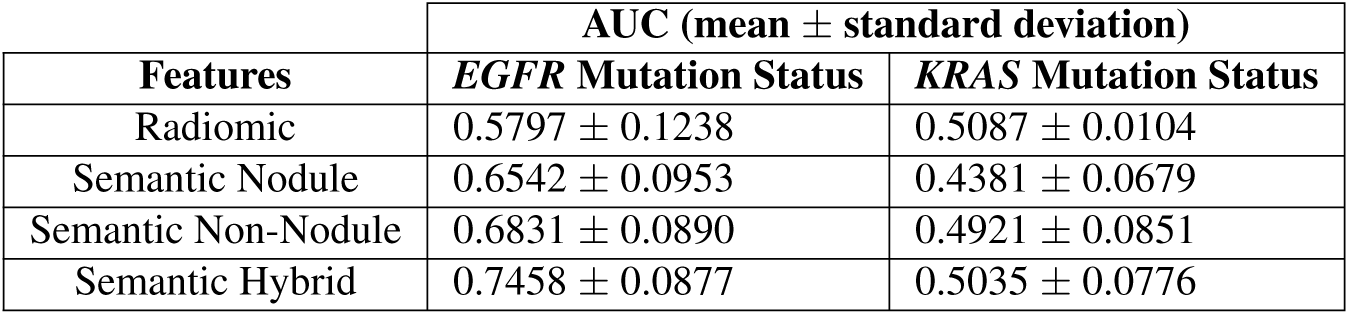
Classification results for *EGFR* and *KRAS* mutation status predictive models considering different sets of input features.

| Features | AUC (mean ± standard deviation) | |
| --- | --- | --- |
|  | *EGFR* Mutation Status | *KRAS* Mutation Status |
| Radiomic | $0.5797 \pm 0.1238$ | $0.5087 \pm 0.0104$ |
| Semantic Nodule | $0.6542 \pm 0.0953$ | $0.4381 \pm 0.0679$ |
| Semantic Non-Nodule | $0.6831 \pm 0.0890$ | $0.4921 \pm 0.0851$ |
| Semantic Hybrid | $0.7458 \pm 0.0877$ | $0.5035 \pm 0.0776$ |

**Figure 2.**
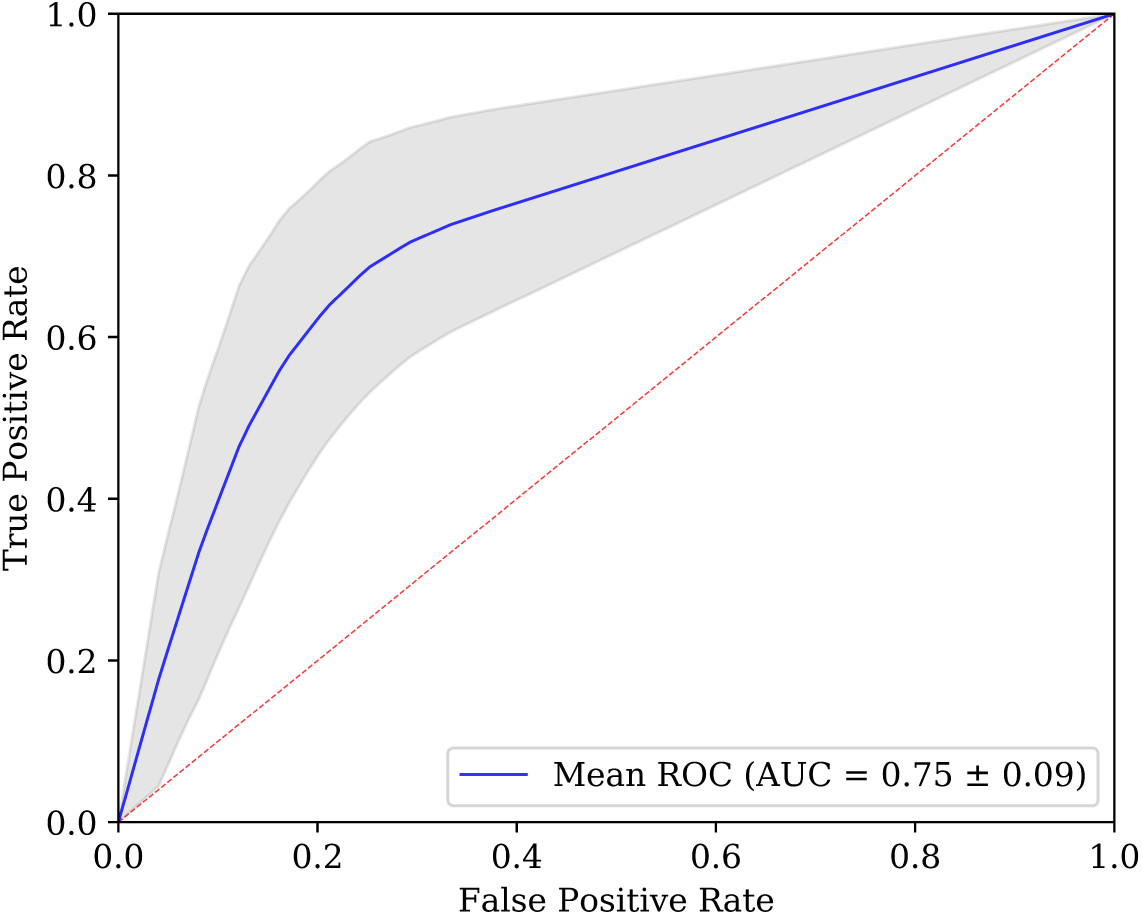
Averaged ROC curve obtained for *EGFR* predictive model based on semantic features. For each of the N=100 runs, the ROC curve is calculated. The blue line depicts the arithmetic average ROC curve and the shading the standard deviation. The red dashed lines indicate ROC curves of at-chance classifiers.

### Most Relevant Features

A subset of features, ranked by importance for the most successful model (*EGFR* mutation status prediction using hybrid semantic features), is presented in Figure 3. They were selected using a minimum threshold of 0.02 and add up to cumulative importance of 0.92 out of 1. The complete list of features importance can be seen in the Supplementary Table 3.

**Figure 3.**
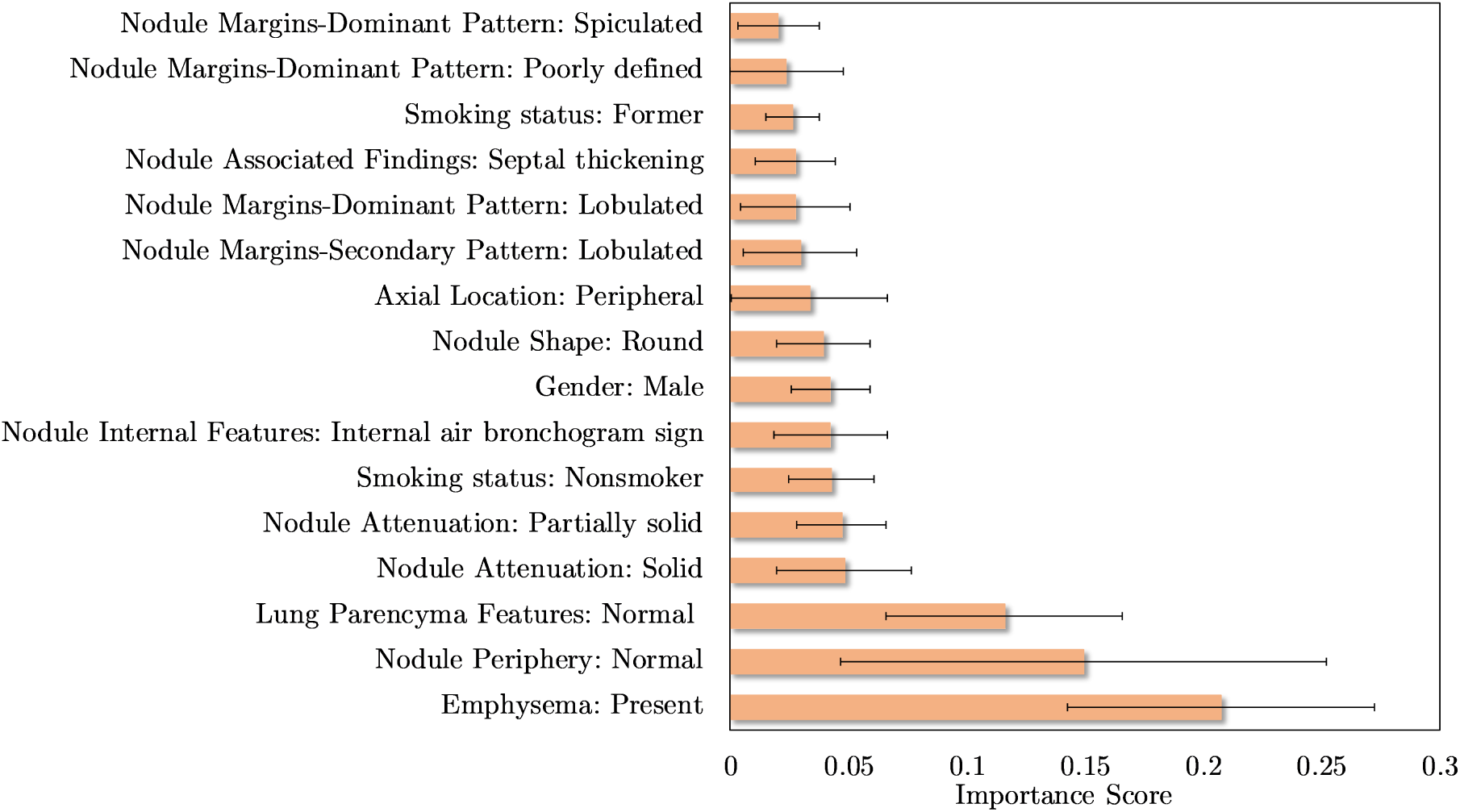
Top 16 semantic features based on the importance scores of features, measured via XGBoost, for predicting the *EGFR* mutation status. Were represented the features that have an average importance score greater than a 0.02. For each of the N=100 runs, the importance score is determined and the average and standard deviation is displayed in the bar graph.

## Discussion and Future Work

The results of the present study suggest that even though *EGFR* mutation status is correlated to CT scans imaging phenotypes, the same does not hold true for *KRAS* mutation status. We hypothesise that this might be due to two reasons: mutated and wild type *KRAS* display identical imaging phenotypes, which is supported by the literature^19, 28, 29^, or our number of samples was too small and unrepresentative to find a relevant pattern for such a complex problem.

The outcomes of this work also indicate that general lung semantic features in conjunction with tumour specific semantic features should be used in order to obtain the best possible *EGFR* mutation status classification results. Only average results were obtained using semantic features that solely describe aspects external to the nodule. The worst performances come from models that only use tumour-describing set of features, either of the radiomic or semantic type. This, combined with the fact that the most relevant features (as determined by the classifier) were tumour external (Figure 1), might hint towards the importance of a holistic lung analysis, instead of a local nodule analysis. Although no previous works profoundly discuss or highlight this particular implication of *EGFR* mutation in imaging phenotypes, there are experiments in the literature that agree with this statement. For example, previous works already showed the importance of extra-tumoral features to obtain a successful *EGFR* mutation status classifier^17, 19^. The most recent review and meta-analysis of CT and clinical characteristics to predict the risk of *EGFR* mutation confirms that CT features with the highest correlation with *EGFR* mutation are from the nodule and other structures of the lung^30^. Also, another work based on deep learning techniques with an interpretable visual output, identified that the regions surrounding the nodule were the most relevant for the classification decision^18, 31^. In our opinion, it is crucial to emphasise this characteristic, as it might change the direction and broaden the analysis spectrum of future radiogenomics studies, which until now have been mainly focusing on the nodule or in a region of interest (ROI) around it^32–34^. Lung cancer is the result of multiple and complex combinations of morphological, molecular and genetic alterations^35^. Since there is a large spectrum of clinicopathological processes that occur during the lung cancer development, it is only natural that important information for the predictive models can be obtained from a larger region of analysis that contains other structures from the lung.

The biggest limitation of this work is the reduced size of the used dataset, which hardly is a good representation of the population affected by lung cancer. In order to better understand the variance in the data and to ensure that the outcomes are not highly influenced by a small number of cases, we used 100 random combinations of cases as train and test sets, reporting the mean values and the standard deviations for the conducted experiences. This limitation is common to the studies in this field, since in general datasets are small and based on patients cohorts from only one medical centre. A reproducible and clinically viable predictive model needs a large and heterogeneous cohort of patients and methods capable of coping with the inherent data heterogeneity^14^. So, a reliable model would require a reliable dataset, collected from multiple centres in order to capture the heterogeneity of the population, but under an uniform protocol to avoid any inconsistency during data record. The access to the data and the uniform acquisition represent the main limitation to build a large dataset. Different protocols used for the data acquisition restrain the mixture of data from different clinical institutions. Additionally, because of privacy issues, the clinical and imaging data is extremely difficult to obtain and requires a large amount of time and effort to submit the protocol to the ethical committees and get approval. Finally, there are other indirect barriers to access the data such as fees and data management requirements^36^. Another limitation of the current study is the number of genes taken into account. The two most frequent gene mutations with lung cancer were selected, however other genes were significantly frequent in this type of cancer and their study could play an important role for novel target therapies, even more personalised and effective.

In the future, new radiomic features should be extracted from radiological images and explored according to this study results. That is, features that reflect the state of pulmonary structures external to the tumour nodule. This would allow us to have a more complete, objective and automatic outlook on the lung, probably delivering more accurate and robust classifiers for *EGFR* mutation status prediction.

## Materials and Methods

### Dataset

The NSCLC-Radiogenomics dataset^13, 21, 22^ comprises data collected between 2008 and 2012 from a cohort of 211 patients with NSCLC referred for surgical treatment, being the only public dataset which comprehends information regarding the mutation status of lung cancer-related genes (*EGFR*, *KRAS* and *ALK*). It contains a set of Computed Tomography (CT) images stored in DICOM format. Since the samples were retrospectively collected, the scanning protocol and scanning parameters were not standardised, thus slice thickness varied from 0.625 mm to 3 mm (median:1.5 mm) and the X-ray current from 124 mA to 699 mA (mean: 220 mA) at 80-140 kVp (mean: 120 kVp). The subjects were in supine position with their arms to the side, while the scans were acquired from the top of the lung to the adrenal gland during a single breath^13^. The nodules segmentation masks are stored as DICOM Segmentation Objects^37^ and are represented as 3D binary images, where voxels belonging to the tumour ROI are represented by the value 1 and voxels outside the tumour ROI are represented by 0. In the cases where the segmentation mask images did not have the same dimensions as their corresponding CT images, the appropriate number of slices was added to the segmentation mask.

Despite including a cohort of 211 NSCLC subjects, only 116 (wild type: 93, mutant: 23) were further considered in the presented radiomic study for *EGFR* mutation status prediction and 114 (wild type: 88, mutant: 26) for *KRAS* mutation status prediction. The scarce availability of tumour masks and target labels did not allow all subjects to be used. Also, Anaplastic lymphoma kinase (ALK), which is the third most frequent oncogene mutated in lung cancer^6^, was not targeted by this study, as the prevalence of mutated cases was too small (wild type: 108, mutant: 2). Additionaly, the dataset comprises a set of subjects whose tumour was analysed by radiologists using 30 nodule and parenchymal features, which describe nodule’s geometry, location, internal features and other related findings. From these subjects, 158 are characterised in terms of *EGFR* mutation status and 157 subjects characterised in terms of *KRAS* mutation status, which were the samples selected for the presented semantic study.

#### Clinical Features

Clinical features were added to the radiomic features as well as to the semantic features to build the predictive models. From now on, we only mention the name of the main group of features that contribute to the models, i.e., radiomic and semantic features. Supplementary Table 1 shows detailed information regarding the clinical data distribution and nomenclature.

#### Radiomic Features

There are image properties, such as the distance between slices, which may differ from scan to scan, and consequently affect the features extracted and the learning ability of the algorithms. Therefore, before trying to extract patterns, the images went through a preprocessing step in order to standardise the scans across the whole dataset.

Firstly, the CT image values were converted to Hounsfield Unites (HU), which is a measure of radiodensity. The HU scale considers the radiodensity of water, at standard temperature and pressure, is defined as 0 HU, and the radiodensity of air is defined as −1000 HU. The equation for computing the HU values based on radiodensity is shown in Equation 1, where *µ* represents the original linear attenuation coefficient of substance, *µ*_*water*_ represents the linear attenuation coefficient of water and *µ*_*air*_ the linear attenuation of air^38^.

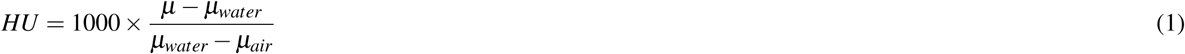

By default, the values returned by the CT scanner are not in this unit. In such manner, the radiodensity values were converted to HU units, by multiplying the voxel value by the slope and adding the intercept related to the linear transformation, which values are stored in the metadata of the scans.

With the purpose of learning patterns from the data using an automatic analysis methodology, it is extremely important that a pixel is represented in the same way in the entire dataset. Having this in mind, the entire dataset (including the tumour masks) were resampled so that neighbour slices and adjacent pixels are separated by 1 mm in *x*, *y* and *z* directions. Images were normalised between −1000 HU and 400 HU, since −1000 HU is the radiodensity of air and values above 400 HU represent hard tissues, not relevant for the task at hand^39^. Values under −1000 HU or above 400 HU were defined as −1000 HU and 400 HU, respectively.

From the 3D images of the nodules of the pre-processed CT scans, a set of 1218 radiomic features were extracted using the open-source package *Pyradiomics*^40^. Features were computed both on the original image and on images obtained after application of wavelet and Laplacian of Gaussian (LoG) filters. A wavelet transform decouples textural information by decomposing the original image in low and high frequencies. A 3D undecimated wavelet transform was applied to each CT image, which decomposed the original image into 8 different images. Considering that *L* is a low-pass filter and *H* a high-pass filter, the original image *X* is decomposed into 8 new images after the wavelet decomposition: *X*_*LLL*_, *X*_*LLH*_, *X*_*LHH*_, *X*_*LHL*_, *X*_*HHH*_, *X*_*HHL*_, *X*_*HLL*_, *X*_*HLH*_. For instance, *X*_*LHL*_ is obtained after applying a low-pass filter along the *x*-dimension, a high-pass filter along the *y*-dimension and a low-pass filter along the *z*-dimension. The remaining images are built in a similar way, applying their respective sequence of low or high-pass filters in *x*, *y* and *z*-direction^41^. Concerning the LoG, five filters with different sigma values were applied (sigma=1.0 mm, 2.0 mm, 3.0 mm, 4.0 mm, 5.0 mm), with the intention of improving texture analysis by detection of multi-scale edges and ridges^42^. In summary, considering the original image and the resulting images after filter application, there were 14 different images to extract features for each sample.

Six classes of features were extracted from the *Pyradiomics* package: shape-based features (14 features), first-order features (18 features), GLCM features (22 features), GLRLM features (16 features), GLSZM features (16 features) and GLDM features (14 features). Shape features include different descriptors of the size and shape of the ROI. Having in mind that shape descriptors are independent of intensity values, only the tumour segmentation masks were required for its computation. First-order features are related to the voxel intensities within the ROI using basic metrics (e.g. mean and standard deviation). GLCM features describe the second-order joint probability function of the ROI. GLRLM features define the length of successive pixels that share the same grey level intensity. GLSZM features quantify the grey level zones in an image, which represent the number of connected voxels that have the same grey level value. At last, GLDM features quantify grey level dependencies in an image. A straightforward overview of the steps involved with the feature extraction is presented in Figure 4.

**Figure 4.**
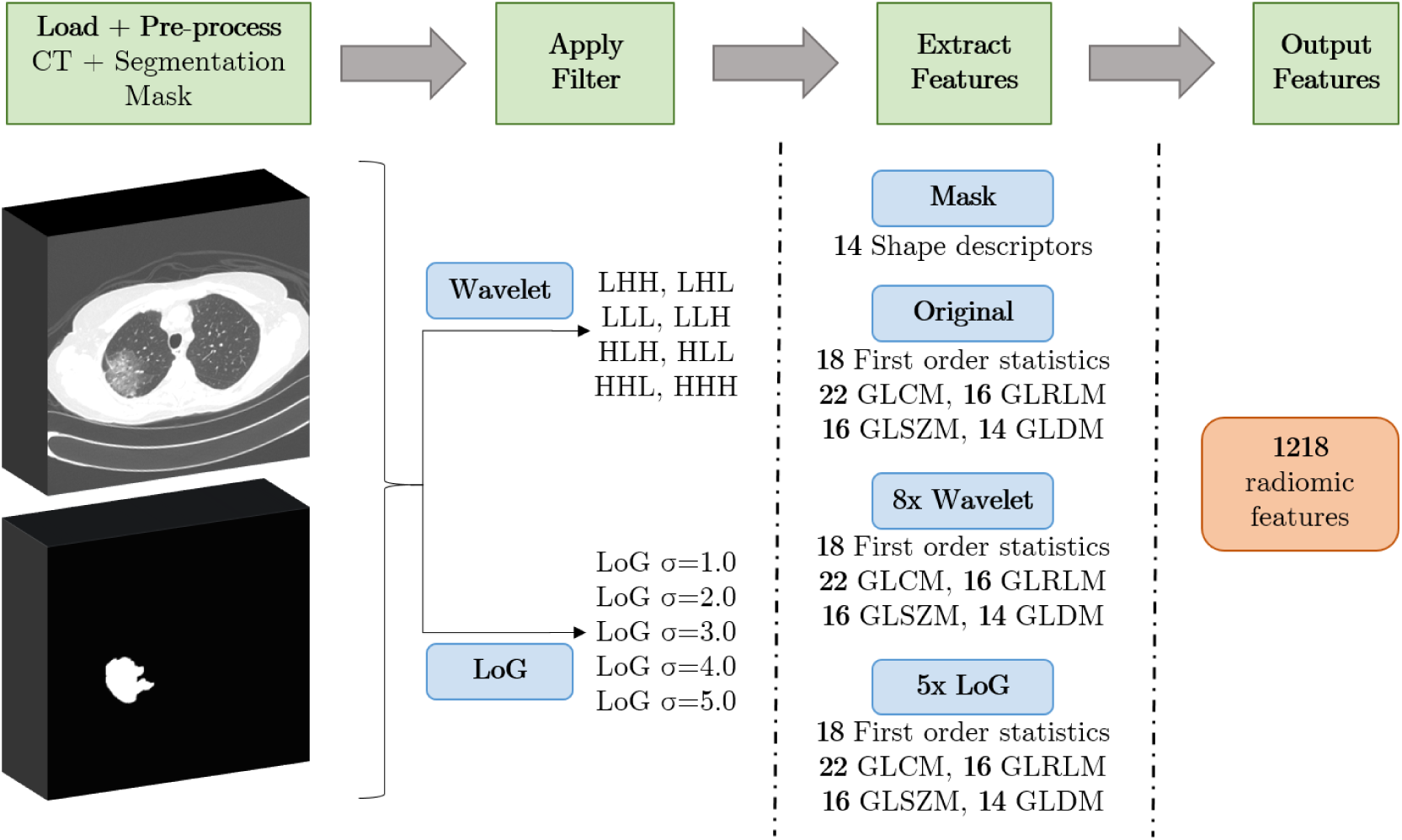
Overview of the process of feature extraction via *Pyradiomics*. First, medical images and segmentation masks are loaded into the software, this step allows to select the region of tumour. Then, after filters have been applied to the original image, radiomic features are extracted from the ROI of the resultant images.

#### Semantic Features

The semantic terms that were used to characterise the patients are common in radiology clinical practice and derive from descriptions in the radiology literature^13^. Definitions of some of the terms used in this description can be found in^43^. The template of semantic terms was developed by two academic thoracic radiologists exclusively for tumours with identifiable nodules and excluded cases without this manifestation. More information about the semantic annotations protocol can be found at^13^.

From the original set of semantic features, some were discarded due to a large number of not applicable values (e.g. the fibrosis type field in a patient that has fibrosis absent), thus, only 18 features were used in the final study. The final dataset comprises percentages of 26% and 25% mutated cases for *EGFR* and *KRAS*, respectively. Supplementary Table 2 shows detailed information regarding the semantic data distribution and nomenclature. Before feeding the data into the model, features were binarised following a one-hot encoding strategy. After that, the number of features increased from 20 to 73.

#### Feature Engineering and Selection

Considering semantic features, most *Lung Parenchyma* categories are underrepresented (see Supplementary Table 1 and 2), with *Normal* and *Bronchial wall thickening* making up to 79.1% and 77.2% of the present *Lung Parenchyma* categories in the *EGFR* and *KRAS* datasets, respectively. To balance the occurrences, we binarise this feature, putting the *Normal* category in a group and the remaining in another, creating a new category titled *Not normal*.

Both semantic and radiomic features were submitted through a process of feature selection, where the correlation matrix was computed and a correlation threshold of 0.95 between variables was set. Additionally, the lower importance radiomic features were excluded. This was done by taking the feature importances from a gradient boosting machine algorithm and only keeping the ones necessary to achieve a cumulative importance of 0.95.

### Dimensionality Reduction

We use Principal Component Analysis (PCA)^44^ followed by t-Distributed Stochastic Neighbour Embedding (t-SNE)^45^ to reduce our high-dimensional data to a two-dimensional space, in order to investigate the existence of class separation between *EGFR* and *KRAS* wild type and mutated samples. PCA allows to find the minimum number of variables that minimise information loss from the original data. This is done by creating new uncorrelated variables (principal components) that maximise variance, which comes down to solving an eigenvector problem. t-SNE is used to further reduce the data dimension to a 2D space. In order to reduce the data dimension, this method minimises the divergence between pairs of input samples (high-dimensional space) and pairs of the corresponding points in the embedding (low-dimensional space) using a cost function.

### Balancing Training Set

In general, machine learning algorithms assume a similar distribution of classes. Here, *EGFR* wild type is over-represented, which could result in a model biased towards this class. However, in this study, the correct classification of both classes is equally important, as the classification of a patient with the wrong mutation status could lead to the administration of a less suitable treatment and, consequently, to shorter progression-free survival. To overcome this class imbalance, Synthetic Minority Over-sampling Technique - Nominal and Continuous (SMOTE-NC) was applied, an extended version of SMOTE generalised to handle data with continuous and nominal features^46^. This technique creates new random synthetic minority class instances between the lines that connect each one of the *n* nearest neighbours of each minority class sample. In comparison to traditional over-sampling, SMOTE-NC has the advantage of building a more general decision region of the minority class. After this algorithm is applied, the training set contains the same number of mutated and wild type samples.

### Classification and Feature Importance

The classifier used in this work was Extreme Gradient Boosting (XGBoost), which is a scalable and accurate implementation of gradient boosted trees algorithms^47^ that has been used for lung cancer related works^48, 49^. A benefit of using gradient boosting is that after the boosted trees are constructed, it is possible to retrieve the importance scores for each feature, based on how useful or valuable each feature was in the construction of the boosted decision trees within the model. Importance is calculated for a single decision tree by the amount that each attribute split point improves the performance measure, weighted by the number of observations the node is responsible for. The feature importance are then averaged across all of the the decision trees within the model.

### Training and Performance Metrics

The training and testing processes were repeated for 100 random splits of the original dataset. Each split comprised a training and test set consisting of 80% and 20% of the original data, respectively. The mean and standard deviation for all the 100 results were reported in favour of reliability and to demonstrate the variance in the data. The classifier hyper-parameters were tuned through a 5-fold cross-validated randomised search on the training data, maximising the models F-measure. The data is balanced individually for each fold using SMOTE-NC, avoiding data leakage. After parameter optimisation, probabilistic outputs of each model with optimal parameters were analysed using the AUC of Receiver Operating Characteristic (ROC). ROC is a probability curve and AUC represents degree or measure of separability, telling how much model is capable of distinguishing between classes. The ROC curve is plotted with True Positive Rate (TPR) against the False Positive Rate (FPR), usually with TPR on the y-axis and FPR on the x-axis.

### Experimental Design

We designed four experiments in order to test and compare which type of input features allow to achieve better performance in gene mutation status prediction. We first trained a model that took nodule-related radiomic features as input. Then, for direct comparison purposes and to allow a modular evaluation, we split the semantic data into three parts: *nodule*, *non-nodule* and *hybrid*. The first one contains only nodular information, the second one contains only information external to the nodule and the third one is the combination of both. The split can be seen in detail in Table 2 of the supplementary material.

## Data Availability

The data was obtained from the open-access NSCLC-Radiogenomics dataset available at the cancer imaging archive (TCIA) database ^13, 21, 22^.

## Additional information

### Competing interests

The authors declare that they have no competing interests.

## Author contributions statement

G.P., T.P., C.D., A.C. and H.O. conceived the experiments, G.P., T.P. and C.D. conducted the experiments, statistical analyses, and manuscript writing. G.P., T.P., J.C., C.F., V.H., A.C. and H.O. performed the clinical interpretation of the results. All authors reviewed the manuscript.

